# Reliable cell retention of mammalian suspension cells in microfluidic cultivation chambers

**DOI:** 10.1101/2022.01.05.475060

**Authors:** Julian Schmitz, Birgit Stute, Sarah Täuber, Dietrich Kohlheyer, Eric von Lieres, Alexander Grünberger

## Abstract

We present a new microfluidic trapping concept to retain randomly moving suspension cells inside a cultivation chamber. In comparison to previously published complex multilayer structures, we achieve cell retention by a thin PDMS barrier, which can be easily integrated into various PDMS-based cultivation devices. Cell loss during cultivation is effectively prevented while diffusive media supply is still ensured.

---

Microfluidic cultivation (MC) of cells under highly controllable environmental conditions is a well-established operation in today’s microfluidics^1^. Here, microfluidic single-cell cultivation (MSCC) with its focus on analyzing single-cell behavior represents a specific subcategory to the field^2^. Typically, MSCC of various single-cell organisms can be realized by the application of cultivation chambers, where cells are trapped in designated regions and cultivated as monolayer colonies for distinct analytical investigations^3^. Due to the spatial restrictions of the cultivation chambers, cells are retained inside the desired compartment and can be analyzed by live cell imaging, resulting in a high temporal resolution of single-cell behavior. With this setting, a broad range of cell types, ranging from algae over bacteria to fungi and mammalians, can be cultivated^4–10^. Commonly, cells inside these cultivation chambers are supplied with nutrients by diffusive mass exchange. For this purpose, the cultivation chambers are arranged along supply channels in which laminar flow of cultivation medium prevails^3^. At the same time supply channels do not only serve for nutrient supply but also represent a potential way for cells to escape the cultivation chambers. Therefore, design and dimension of supply channel, cultivation chamber, and the chamber’s entrance always represents a tradeoff between optimal nutrient supply and sufficient cell retention. Especially for the long-term cultivation of slow growing cells^11^ as well as the microfluidic cultivation of motile cells^12^, a reliable cell retention concept is a fundamental requirement to prevent permanent cell loss, which otherwise compromises qualitative and quantitative cell studies.

In the last years, MSCC for mammalian cell lines, in particular for Chinese hamster ovary (CHO) cells, became of increasing interest to investigate growth and robustness of industrially relevant production cell lines^13,14^. Here, especially suspension cells are of industrial relevance, as large-scale bioproduction processes exclusively use suspension cell lines^15,16^. In comparison to bacteria and yeast, which are traditionally kept inside the cultivation chambers by squeezing them tightly into narrow chamber heights, because of their deformable nature CHO cells cannot be retained inside the chamber the same way as cells with rigid cell walls. Although they are not able to move actively, when cultivated in MSCC devices CHO suspension cells also randomly migrate inside the boundaries of the cultivation chambers^13,17^. Unfortunately, these random movements frequently lead to cell loss, as migrating cells leave the chamber through the entrance]. Additionally, squeezing cells inside narrow microfluidic structures potentially influences cellular behavior because of spatial restriction and thereby could lead to compromised growth. Thus, other approaches to reliably retain cells must be established to permit quantitative analysis of single-cell cultivation.

In order to overcome this challenge, a variety of microfluidic devices with different approaches to retain (motile) cells have been published over the years. Several of these setups rely on PDMS multilayer chips to either trap cells at the bottom of a cultivation well^18,19^ or to establish on-chip valving to lock cultivation chambers after cells have been trapped^20–22^. Others exhibit cultivation chambers made of SU8, that are combined with fluid control layers made of PDMS-membrane hybrids to ensure supply of the cultivated cells with medium^23^. Although very sophisticated and efficient, all these approaches are very complex and are prone to malfunction, since their setup depends on a multitude of fabrication steps and technical components.

In this respect several microfluidic designs consisting in a PDMS single-layer chip have been introduced as well. Some rely on dead-end cultivation chambers which are sealed with air after cell loading, to retain cells inside the chamber^24^. While cell retention is highly reliable, environmental conditions in sealed cultivation chambers resemble a batch-mode cultivation and thus are subjected to a drastic environmental changes over long-term cultivation and thereby disqualify these designs for cultivations, where defined environmental control is desired. Other concepts rely on cultivation chambers with narrow entrances, minimizing the cross section between cultivation chamber and supply channels and thereby minimizing the probability of cell loss^25^.

In this work, we developed an easy-to-integrate PDMS barrier for our previously developed microfluidic cultivation device (Fig. 1A)^13^ that enables enhanced cell retention by introducing a physical blocking structure into the cultivation chamber’s entrances (Fig. 1B), that only can be passed by applying pressure during cell loading but does not permit the escape of randomly moving CHO cells during cultivation (Fig. 1C). Here, pressure-induced deformation of cells is assumed to be the decisive factor in cell loading, yet slight bending of the PDMS barrier might additionally promote this process. As can be seen in Fig. 1B, the barrier structure exhibits the same height as the chamber, so that it fully locks up the entrance and does not function as a movable hatch. With a width of 27 μm the barrier nearly closes the whole entrance except for a 1.5 μm wide and 2.0 μm high gap (Fig. 1D). The compatibility of the PDMS barrier with our previously developed MSCC design ensures simple chip fabrication and easy application for the cultivation of CHO suspension cells. Consisting of four parallel supply channels, the device exhibits in total 60 cultivation chambers, 30 of them are arranged in line between two supply channels (Fig. 2A). To ensure monolayer growth of trapped cells, the modified design holds a limited chamber height of approx. 10 μm (Fig. S1A). Based on our previous design, the ratio between supply channel and cultivation chamber height remained constant (approx. 2:1) to restrict laminar flow to the supply channels which results in exclusively diffusive mass exchange between channel and chamber. Likewise, the supply channel width of 200 μm and the cultivation chamber’s base area of 200 x 200 μm^2^ stayed unchanged since it allows MSCC experiments for up to 7 days until cells outgrow the limited space (Fig. S1B&C).

**Fig. 1:**
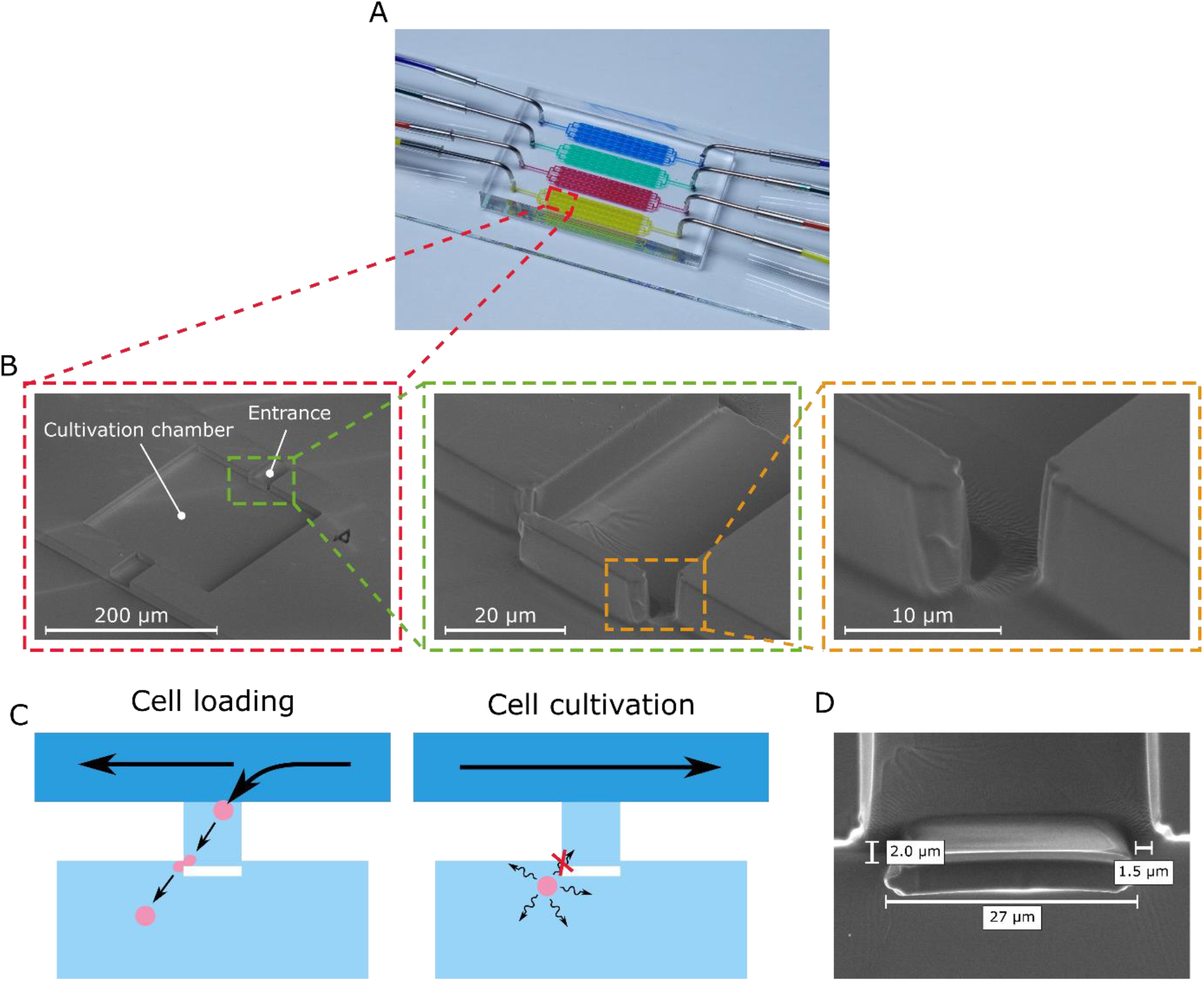
Structure and trapping concept of the MSCC device for CHO suspension cell lines with enhanced cell retention. (A) Microfluidic PDMS-glass-based cultivation device (B) Scanning electron microscopy image of the microfluidic structure illustrating the devices dimensions and trapping barrier. (C) Schematic drawing of the cell trapping concept based on a PDMS barrier that is traversable by applying pressure during cell loading but non-traversable by random cellular movement during cultivation. (D) Scanning electron microscopy image of the PDMS barrier located in the cultivation chamber’s entrance.

**Fig. 2:**
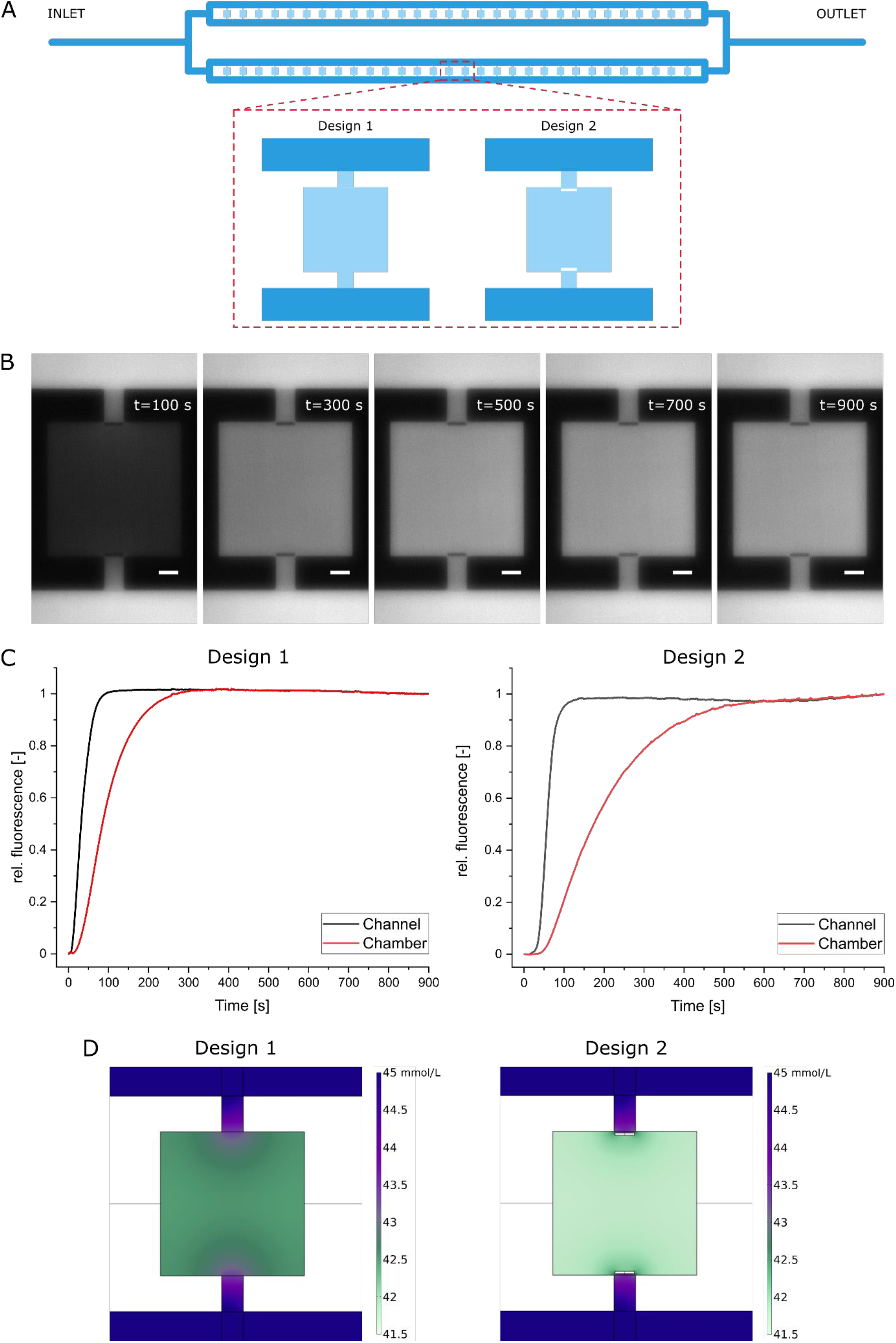
Microfluidic characterization of the MSCC designs. (A) Individual cultivation array that contains 60 cultivation chambers with either our previous chamber design (Design 1)^13^ or the chamber design from this work (Design 2). (B) Image sequence of trace substance experiments to quantify diffusive mass exchange for the MSCC device with design 2. Scale bar=30 μm. (C) Medium exchange duration until full equilibrium between channel and chamber is achieved based on rel. fluorescein signal for both designs. (D) Glucose concentration profile during MSCC cultivation assuming a steady state with 181 cells inside the chamber with a constant glucose uptake rate of 3800 nmol per 10^6^ cells and day for both designs.

Nearly closing the whole entrance of the cultivation chamber makes seeding cells more difficult. Therefore, we applied air to the microfluidic cultivation device to create a directed flow through the cultivation chambers (Fig. S2). However, this procedure must be conducted very carefully, otherwise cells will be sheared when passing through the narrowing between barrier and cultivation chamber wall. Additionally, no air bubble must remain inside the microfluidic cultivation device during the subsequent perfusion, or the intended flow profile will be disturbed.

As a result from almost closing the cultivation chamber’s entrance, not only loading characteristics of our new microfluidic design are drastically changed, but also mass exchange between channel and chamber is decreased. During MSCC experiments environmental conditions relating to nutrient concentrations are kept constant due to steady perfusion of the cultivation device. However, as cells start to fill the chamber, limitations might occur when cellular uptake rate outruns diffusive mass exchange. Therefore, we quantified diffusive mass exchange as well as glucose concentration profiles inside the presented device by fluorescein trace substance experiments and computational fluid dynamics (CFD) simulations according to the model published by Schmitz et al. 2020^13^ for our already established design (Design 1) and our newly developed design (Design 2) with enhanced cell retention capability.

Fig. 2B and the respective video (Video S1) show that the fluorescein signal inside the cultivation chamber of design 2 increases clearly delayed to the fluorescence inside of the supply channels. As can be seen from the rel. fluorescence level on Fig. 2C it takes 600 s until the same medium conditions from the supply channels are present inside the cultivation chamber. In comparison to our previously published MSCC device (design 1), where equilibrium was reached after approx. 300 s (Video S2, Fig. 2C), this is a delay of 100 %. However, since our device is operated in a static way with constant cultivation conditions, we do not expect any problems caused by the prolonged medium exchange duration, since chambers are inoculated with only few cells and cellular growth (t_D_=15 h) is slow in comparison to the determined medium exchange duration (t_exchange_=600 s). Additionally, the CFD simulation indicates that with approx. 180 cells inside a single cultivation chamber, which is a representative cell number per chamber after 150 h of cultivation, no glucose limitation occurs, given that the minimal glucose concentration does not drop below 41.5 mmol/L (Fig. 2c).

Multiple cultivation experiments with both cultivation chamber designs were performed to evaluate the cell retention capacity of the novel trapping concept against the basic design. By quantifying the cell number during cultivation, we recorded the growth of three microcolonies in both designs. Fig. 3A shows the growth progression of three microcolonies that were cultivated using the novel trapping concept. As the starting cell number varies between one and three cells, the rise of the curves is slightly delayed in time but, as Fig. 3B clearly indicates, strictly exponential and the specific growth rates of the respective microcolonies match. In contrast to that, the growth curves of the three microcolonies cultivated with the basic trapping concept feature distinct bends, which correlate with the loss of cells from the cultivation chamber (Fig. 3C). The related semi-logarithmic plot illustrates the influence of losing cells on the specific growth rate, which is clearly decreased when determined over the whole cultivation time (Fig. 3D). This difference in cell retention and its influence on microcolony growth becomes apparent when looking at the corresponding videos (Video S3 & S4). With design 2 cells only leave the chamber when they are pushed out by other cells or directly divide throughout the narrow gap between barrier and chamber wall.

**Fig. 3:**
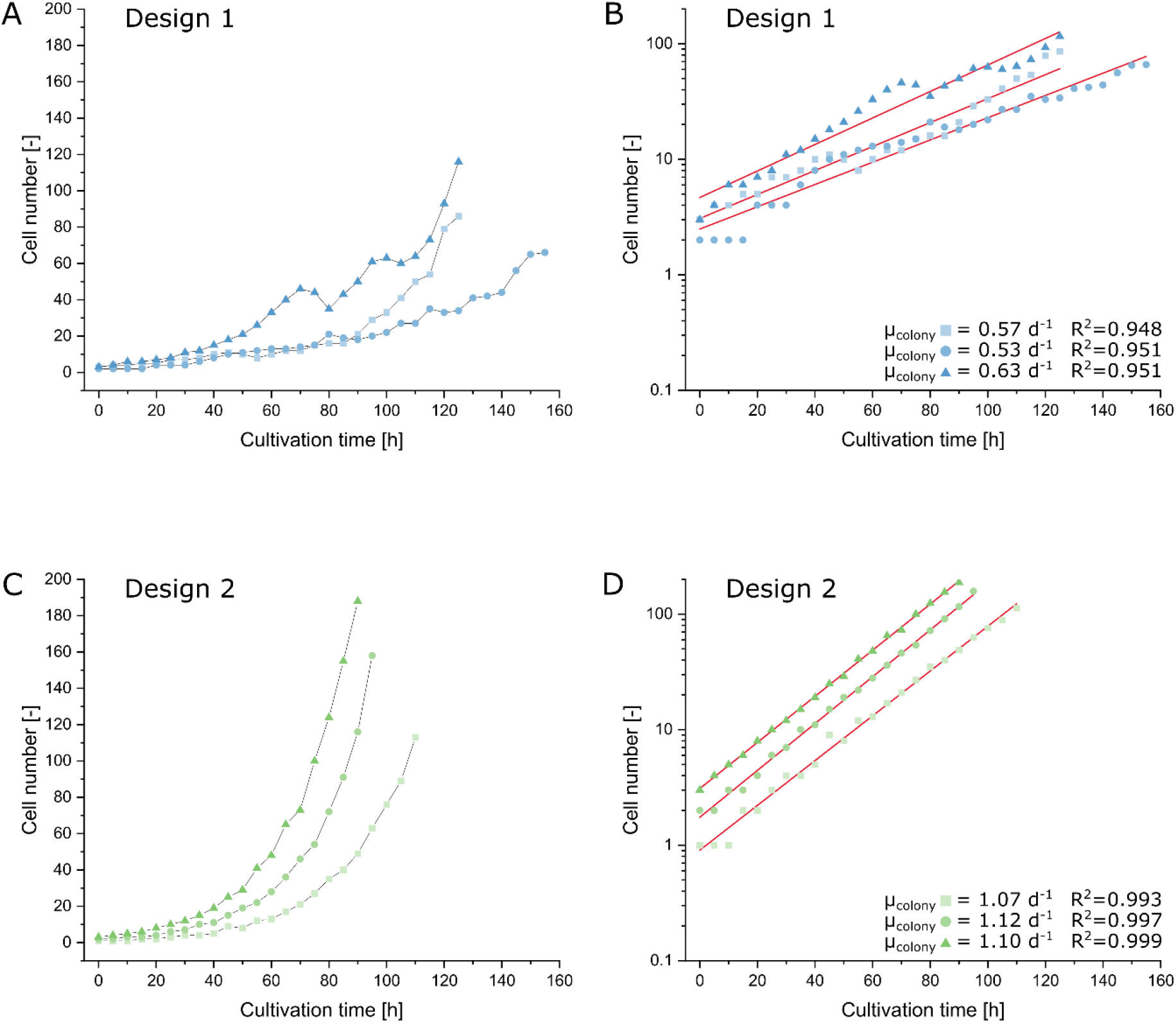
MSCC of CHO cells and cell retention assessment of the MSCC designs. (A) Growth of three CHO-K1 microcolonies cultivated applying design 1. (B) Semi-logarithmically plotted growth profile of the microcolonies cultivated applying design 1. (C) Growth of three CHO-K1 microcolonies cultivated applying design 2. (D) Semi-logarithmically plotted growth profile of the microcolonies cultivated applying design 2.

Cell loss during long-term cultivation leads to a distinct underestimation of specific growth rates μ when determined on colony level and thereby results in immense differences between growth rate estimation on single-cell level μ_single-cell_ and on colony level μ_colony_ ^13^. In order to have a closer look on this discrepancy, we additionally analyzed growth on the single-cell level by determining the doubling time t_D_ of cellular division events during the cultivation. Here, the comparison between cells cultivated with both designs shows no significant difference (Fig. 4).

**Fig. 4:**
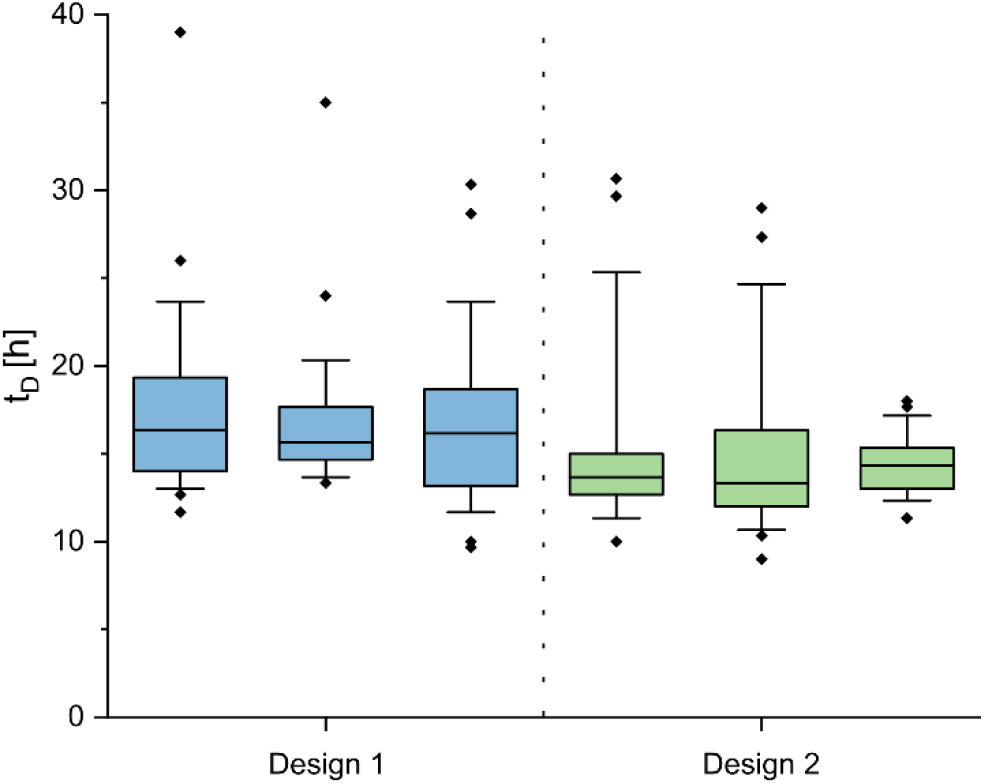
Comparison of single-cell division behavior between the two microfluidic trapping concepts. Depicted are the singlecell doubling times t_D_ of cells cultivated in chambers with both designs. The colored segment marks the interquartile range from 25% to 75%, the horizontal lines show the median. The whiskers represent the 10% and 90% percentile and the tilted squares mark rare cellular events.

By assuming exponential growth, t_D_ can be converted into μ applying equation (1):

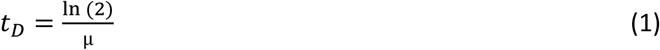

Comparing the above determined μ_colony_ with the average μ_single-cell_ of the same microcolonies, it is clearly noticeable that these two values show only small variations when cells are cultivated in the cultivation chambers design 2. However, cultivating cells under reoccurring cell loss with design 1 leads to a significant difference (Tab. 2).

**Tab. 1:** Overview of recently published microfluidic cultivation devices with enhanced cell retention. The listed approaches are itemized concerning the characteristics of the respective cultivation area, the cell retention method, and the nutrient supply. Additionally, the cultivated cell type is indicated.

| Cultivation area | Retention method | Nutrient supply | Cultivated cell type | Author |
| --- | --- | --- | --- | --- |
| Wells | Sedimentation | Diffusive | Mouse hematopoietic cells | Lecault et al.<sup>18</sup> |
| Wells | Sedimentation | Diffusive | Mouse ovarian cancer cells | Dadgar et al.<sup>19</sup> |
| Chambers | Quake valves | Convective | *Chlamydomonas reinhardtii* | Eu et al.<sup>20</sup> |
| Chambers | Quake valves | Diffusive | Primary murine cells, Murine embryonic stem cells | Dettinger et al.<sup>21</sup> |
| Chambers | Quake valves | Convective | Human primary mesenchymal stem cells | Gómez-Sjöberg et al.<sup>22</sup> |
| Chambers | Membrane sealing | Diffusive | *Dictyostelium discoideum* | Delince et al.<sup>23</sup> |
| Chambers | Air sealing | None | *Salpingoeca rosetta* | Halperin et al.<sup>24</sup> |
| Chambers | Narrow entrances | Diffusive | Chinese hamster ovary | Kolnik et al.<sup>25</sup> |

**Tab. 2:** Comparison of the single-cell growth rate data μ_single-cell_, calculated from the geometrical mean of the determined single-cell doubling times t_D_, with the colony growth rate data μ _colony_, determined graphically based on the cell number, for both designs.

| | | $\bar{t}_{D, \text{geom}}$ | $\mu_{\text{single-cell}}$ | $\mu_{\text{colony}}$ |
| --- | --- | --- | --- | --- |
| Design 1 | Chamber 1 | 16.88 h | 0.99 h<sup>-1</sup> | 0.57 h<sup>-1</sup> |
|  | Chamber 2 | 16.81 h | 0.99 h<sup>-1</sup> | 0.53 h<sup>-1</sup> |
|  | Chamber 3 | 16.07 h | 1.04 h<sup>-1</sup> | 0.63 h<sup>-1</sup> |
| Design 2 | Chamber 1 | 14.83 h | 1.12 h<sup>-1</sup> | 1.07 h<sup>-1</sup> |
|  | Chamber 2 | 14.52 h | 1.15 h<sup>-1</sup> | 1.12 h<sup>-1</sup> |
|  | Chamber 3 | 14.34 h | 1.16 h<sup>-1</sup> | 1.10 h<sup>-1</sup> |

Owing to the modular structure of our novel trapping concept and its simple way of retaining (motile) cells, the trapping principle can be transferred to other cultivation chamber-based designs and thus be applied for the cultivation of other organisms in general. Furthermore, not only singlecell growth studies might be feasible but also a broad variety of taxis or migration studies in restricted compartments is conceivable^26^. Therefore, we believe that our trapping concept can have a wider field of application than just the cultivation of cells.

## Conclusions

The non-invasive single-cell trapping and cultivation of motile cells in microfluidic devices always represents a difficult challenge. Especially when coupled with live cell imaging, cellular growth and movement must be spatially restricted, otherwise single-cell cultivation and analysis is not feasible due to constant loss of individual cells. As CHO suspension cells show random movement inside a microfluidic cultivation chamber, we tackled the challenge of cell loss by proposing a novel trapping concept based on the introduction of a physical blocking structure into a previously published MSCC device^13^. During cell loading the barrier can be passed by cells, afterwards the barrier prevents cells from escaping the cultivation chamber. At the same time, single cells can be cultivated for multiple days without any nutrient limitations, although diffusive cross section between supply channel and cultivation chamber was decreased.

By introducing our novel trapping concept to the field of microfluidic single-cell cultivation, systematic studies of cellular behavior can be performed in a reproducible way without decreased significance due to constant loss of analyzed cells. Yet, we believe that our cell retention approach will not only proof valuable in the field of bioprocess microfluidics but will also pave the way for future cellular migration assays in the context of basic or biomedical single-cell research.

## Author Contributions

Conceptualization: JS, AG. Investigation: JS, BS. Formal analysis and validation: JS, ST, AG. Visualization: JS, BS. Resources: DK, EvL, AG. Supervision: JS, EvL, AG. Writing – original draft: JS, AG. Writing – review & editing: JS, AG.

## References

1 A. Grünberger, N. Paczia, C. Probst, G. Schendzielorz, L. Eggeling, S. Noack, W. Wiechert and D. Kohlheyer, A disposable picolitre bioreactor for cultivation and investigation of industrially relevant bacteria on the single cell level, Lab on a chip, 2012, 12, 2060–2068.

2 A. Grünberger, W. Wiechert and D. Kohlheyer, Single-cell microfluidics: opportunity for bioprocess development, Current opinion in biotechnology, 2014, 29, 15–23.

3 A. Grünberger, C. Probst, S. Helfrich, A. Nanda, B. Stute, W. Wiechert, E. von Lieres, K. Nöh, J. Frunzke and D. Kohlheyer, Spatiotemporal microbial single-cell analysis using a high-throughput microfluidics cultivation platform, Cytometry Part A, 2015, 87, 1101–1115.

4 P. J. Graham, J. Riordon and D. Sinton, Microalgae on display: a microfluidic pixel-based irradiance assay for photosynthetic growth, Lab on a chip, 2015, 15, 3116–3124.

5 A. Groisman, C. Lobo, H. Cho, J. K. Campbell, Y. S. Dufour, A. M. Stevens and A. Levchenko, A microfluidic chemostat for experiments with bacterial and yeast cells, Nature methods, 2005, 2, 685–689.

6 A. C. Rowat, J. C. Bird, J. J. Agresti, O. J. Rando and D. A. Weitz, Tracking lineages of single cells in lines using a microfluidic device, Proceedings of the National Academy of Sciences of the United States of America, 2009, 106, 18149–18154.

7 C. S. Luke, J. Selimkhanov, L. Baumgart, S. E. Cohen, S. S. Golden, N. A. Cookson and J. Hasty, A Microfluidic Platform for Long-Term Monitoring of Algae in a Dynamic Environment, ACS synthetic biology, 2016, 5, 8–14.

8 G. Ullman, M. Wallden, E. G. Marklund, A. Mahmutovic, I. Razinkov and J. Elf, High-throughput gene expression analysis at the level of single proteins using a microfluidic turbidostat and automated cell tracking, Philosophical transactions of the Royal Society of London. Series B, Biological sciences, 2013, 368, 20120025.

9 S. Demming, B. Sommer, A. Llobera, D. Rasch, R. Krull and S. Büttgenbach, Disposable parallel poly(dimethylsiloxane) microbioreactor with integrated readout grid for germination screening of Aspergillus ochraceus, Biomicrofluidics, 2011, 5, 14104.

10 H. S. Kim, T. P. Devarenne and A. Han, A high-throughput microfluidic single-cell screening platform capable of selective cell extraction, Lab on a chip, 2015, 15, 2467–2475.

11 K. Ziółkowska, A. Stelmachowska, R. Kwapiszewski, M. Chudy, A. Dybko and Z. Brzózka, Long-term three-dimensional cell culture and anticancer drug activity evaluation in a microfluidic chip, Biosensors & bioelectronics, 2013, 40, 68–74.

12 V. Tokárová, A. Sudalaiyadum Perumal, M. Nayak, H. Shum, O. Kašpar, K. Rajendran, M. Mohammadi, C. Tremblay, E. A. Gaffney, S. Martel and D. V. Nicolau, Patterns of bacterial motility in microfluidics-confining environments, Proceedings of the National Academy of Sciences, 2021, 118. DOI: 10.1073/pnas.2013925118.

13 J. Schmitz, S. Täuber, C. Westerwalbesloh, E. von Lieres, T. Noll and A. Grünberger, Development and application of a cultivation platform for mammalian suspension cell lines with single-cell resolution, Biotechnology and bioengineering, 2021, 118, 992–1005.

14 M. P. Marques and N. Szita, Bioprocess microfluidics: applying microfluidic devices for bioprocessing, Current opinion in chemical engineering, 2017, 18, 61–68.

15 G. Walsh, Biopharmaceutical benchmarks 2018, Nature biotechnology, 2018, 36, 1136–1145.

16 M. M. Zhu, M. Mollet, R. S. Hubert, Y. S. Kyung and G. G. Zhang, eds., Industrial Production of Therapeutic Proteins: Cell Lines, Cell Culture, and Purification.

17 J. Schmitz, O. Hertel, B. Yermakov, T. Noll and A. Grünberger, Growth and eGFP Production of CHO-K1 Suspension Cells Cultivated From Single Cell to Laboratory Scale, Frontiers in bioengineering and biotechnology, 2021, 9, 716343.

18 V. Lecault, M. Vaninsberghe, S. Sekulovic, D. J. H. F. Knapp, S. Wohrer, W. Bowden, F. Viel, T. McLaughlin, A. Jarandehei, M. Miller, D. Falconnet, A. K. White, D. G. Kent, M. R. Copley, F. Taghipour, C. J. Eaves, R. K. Humphries, J. M. Piret and C. L. Hansen, High-throughput analysis of single hematopoietic stem cell proliferation in microfluidic cell culture arrays, Nature methods,2011, 8, 581–586.

19 N. Dadgar, A. M. Gonzalez-Suarez, P. Fattahi, X. Hou, J. S. Weroha, A. Gaspar-Maia, G. Stybayeva and A. Revzin, A microfluidic platform for cultivating ovarian cancer spheroids and testing their responses to chemotherapies, Microsystems & nanoengineering, 2020, 6, 93.

20 Y.-J. Eu, H.-S. Park, D.-P. Kim and J. Wook Hong, A microfluidic perfusion platform for cultivation and screening study of motile microalgal cells, Biomicrofluidics, 2014, 8, 24113.

21 P. Dettinger, T. Frank, M. Etzrodt, N. Ahmed, A. Reimann, C. Trenzinger, D. Loeffler, K. D. Kokkaliaris, T. Schroeder and S. Tay, Automated Microfluidic System for Dynamic Stimulation and Tracking of Single Cells, Anal. Chem., 2018, 90, 10695–10700.

22 R. Gómez-Sjöberg, A. A. Leyrat, D. M. Pirone, C. S. Chen and S. R. Quake, Versatile, fully automated, microfluidic cell culture system, Anal. Chem., 2007, 79, 8557–8563.

23 M. J. Delincé, J.-B. Bureau, A. T. López-Jiménez, P. Cosson, T. Soldati and J. D. McKinney, A microfluidic cell-trapping device for single-cell tracking of host-microbe interactions, Lab on a chip, 2016, 16, 3276–3285.

24 S. O. Halperin, C. T. Poling, S. R. Mathrani, B. W. Turner, A. C. Greene, M. E. Dueck and F. B. Myers, A massively parallel microfluidic device for long-term visualization of isolated motile cells, Microfluid Nanofluid, 2014, 17, 821–829.

25 M. Kolnik, L. S. Tsimring and J. Hasty, Vacuum-assisted cell loading enables shear-free mammalian microfluidic culture, Lab on a chip, 2012, 12, 4732–4737.

26 A. L. Höving, J. Schmitz, K. E. Schmidt, J. F. W. Greiner, C. Knabbe, B. Kaltschmidt, A. Grünberger and C. Kaltschmidt, Human Blood Serum Induces p38-MAPK-and Hsp27-Dependent Migration Dynamics of Adult Human Cardiac Stem Cells: Single-Cell Analysis via a Microfluidic-Based Cultivation Platform, Biology, 2021, 10. DOI: 10.3390/biology10080708

