## Supplementary Information for "Reliable cell retention of mammalian suspension cells in microfluidic cultivation chambers"

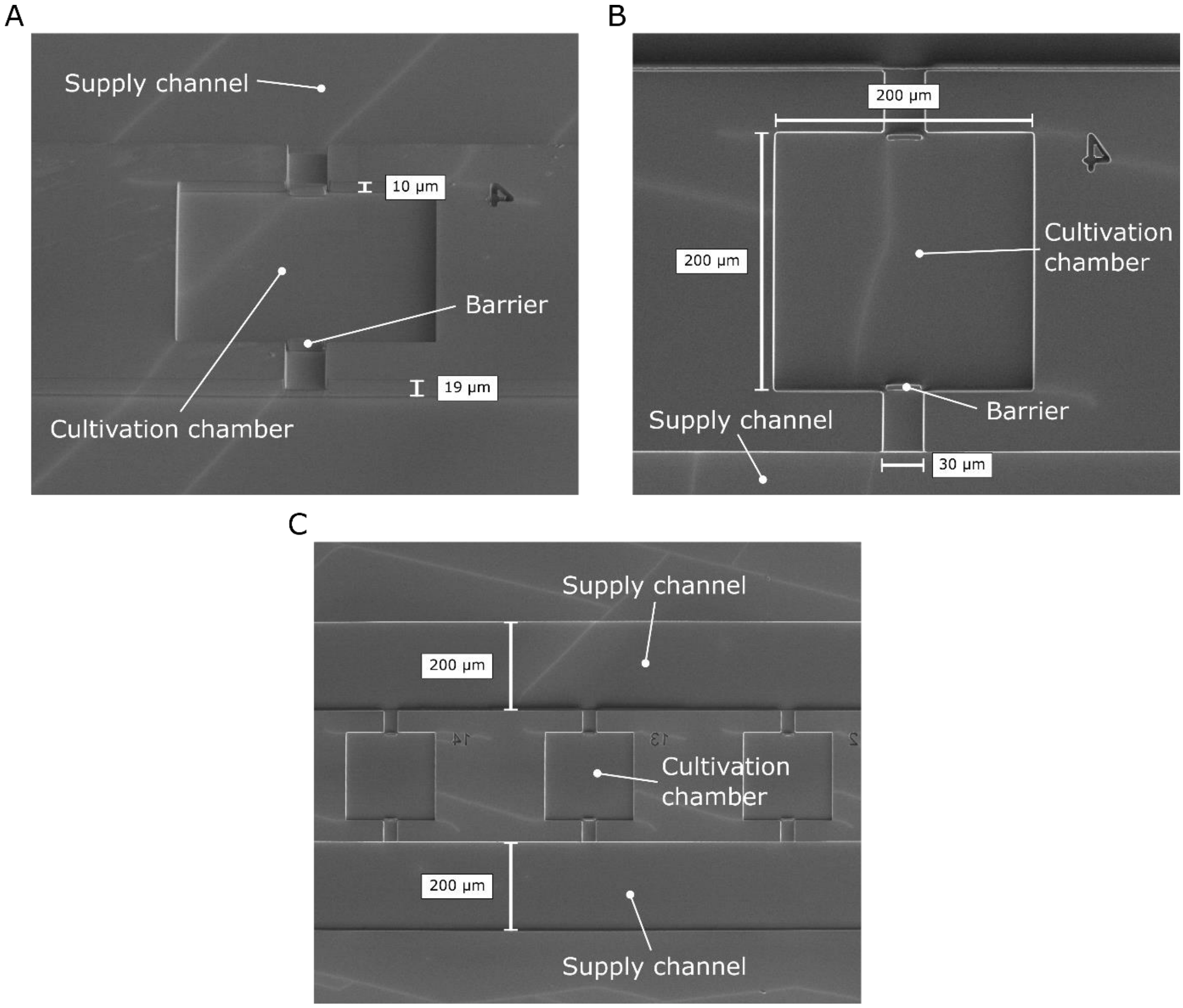


Fig. S1: Scanning electron microscope (SEM) images of the microfluidic cultivation device with PDMS barrier in the cultivation chamber's entrance. (A) The approx. height of the cultivation chamber is 10 µm while the supply channel is approx. 19 µm high. (B) The base area of the cultivation chamber is 200 x 200 $µm^2$ and the chamber entrance has a width of 30 µm. (C) The displayed image section shows three cultivation chambers that are lined up along the adjacent supply channels. The channels have a width of 200 µm.

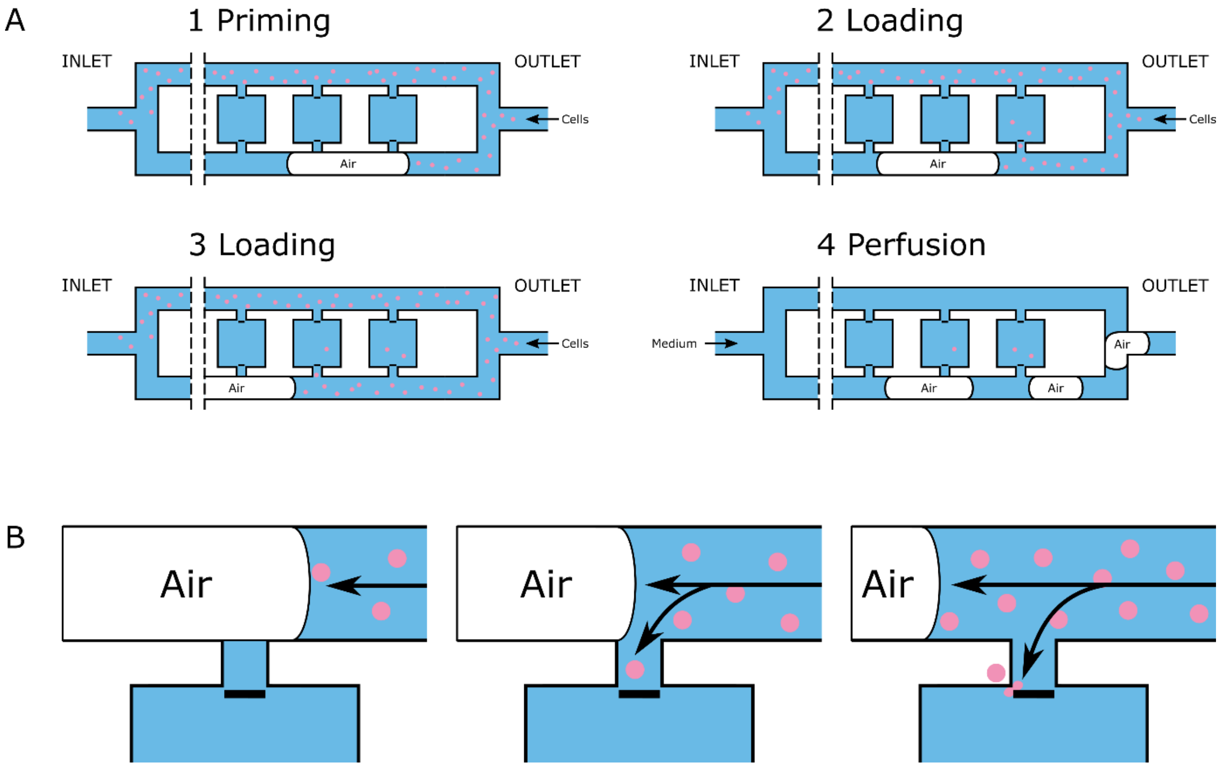


Fig. S2: Loading procedure to capture cells inside the MSCC device with the novel trapping concept. (A) The MSCC device is manually flushed with cell suspension from the outlet side using a single-use syringe. By introducing air into the supply channels, one adjacent channel is blocked so that the cultivation chambers can directly be flushed with cell suspension. After sufficient loading, medium is pumped through the MSCC device from the inlet side so that remaining air is pushed out of the channels and constant perfusion can be established. (B) Zoom-in of the chamber's entrance. Once the entrance is no longer blocked by air, the flow is directed into the cultivation chamber. Due to the increased pressure of manual loading, cells are pushed through the narrow gap between barrier and the walls of the respective cultivation chamber.
